# Immersion as inference: An active inference account of induced realities and its relation to the two-competitor model

**DOI:** 10.64898/2026.11.03.764079

**Authors:** Laura Renz, Kyra L. GauthierDickey, Jonas Vibell, Alexander L. Francis, Farah I. Corona-Strauss, Daniel J. Strauss

## Abstract

Immersive technologies are designed for a single target: the human perceptual system. Engineers of virtual and augmented reality have long recognized that a display does not have to replicate the physical world, it only has to provide the inputs that satisfy the senses at the level at which the brain samples them. This engineering intuition is, in fact, a statement about brain function. Here we develop it into a formal account of immersion within the active inference framework. We treat an induced reality (a virtual, augmented, or mixed reality, but also a book or a daydream) as a candidate generative model that competes with the model of the actual reality for the explanation of a shared sensorium and for the control of action. Immersion is then defined as the ensuant belief that the induced reality best explains the current sensory signals and supports the current perceived reality of the winning model, despite any break in immersion by Bayesian model switching driven by a high-precision prediction error that the induced model cannot absorb. Exogenous and endogenous attention, implemented as weights governing stream selection in our earlier two-competitor model, which treated attention as necessary but not sufficient for immersion, appear as two routes of precision allocation: likelihood precision of evolutionarily entrenched sensory events, and precision that follows the expected free energy of policies. Affect and motivation enter through the pragmatic and epistemic value of remaining in the induced reality and through the rate of change of free energy. We show that the two-competitor attentional focus vector and its angle are recovered from the active inference variables, and we reproduce the two scenarios of the original model, a rewarding and a fatiguing activity interrupted by a slamming door, in a minimal simulation. The account links presence, embodiment, cybersickness, and narrative absorption to a single quantity, and it yields concrete predictions for the psychophysiological assessment and closed-loop control of immersion. We discuss implications for neuroergonomic design in which the evolutionarily entrenched core of perception is non-negotiable while its experience-dependent periphery can be trained.

## 1 Introduction

Interest in sensory-driven immersion has increased alongside advances in affordable head-mounted displays, spatial audio, haptic interfaces, and the business models built around a digital virtual universe [1]. Yet a broadly accepted definition of immersion in terms of established neurocognitive constructs and measurable quantities has remained elusive. In our earlier work, we approached this problem from the perspective of attention and proposed the two-competitor model [1]. The model treats an induced reality (a virtual, augmented, or mixed reality, but also a novel or a daydream) and the actual reality as two orthogonal dimensions competing for the projection of an attentional focus vector, and it integrates the taxonomy of exogenous versus endogenous attention with the taxonomy of external versus internal attention described by Chun et al. [2]. The angle of the attentional focus vector relative to the actual reality axis quantifies the degree of immersion, and its magnitude quantifies the total allocation of attentional resources. Affect and motivation enter the model through a fatigue module that maps reward and effort onto the endogenous weights [3, 4]. The central hypothesis of that work was that attention is a necessary but not a sufficient condition for immersion: without attention to the induced reality there is no immersion, but attention alone does not produce it, because the stimuli must also resonate with the goals and the affective state of the person. The two-competitor model captured this resonance only through the empirical weights of its fatigue module.

The two-competitor model was deliberately conceptual. It provided a taxonomy and a graphical intuition, and it allowed a quantitative implementation, but it left open many questions that a neuroscience of immersion must eventually answer: Why does an induced reality “win” the competition at all, given that the physical world never disappears from the sensorium? What does it mean, in terms of brain computation, for the stimuli of an induced reality to “fit together in some particularly compelling way” [1]? Why is immersion so fragile win respect to interruptions such as a a slamming door but so robust with respect to e.g. the weight of a headset? How do affect, motivation, and effort modulate immersion beyond the empirical weights that we assigned to them? Why is attention, which is plainly necessary for immersion, not sufficient for it? And how does immersion relate to the sensory conflict thought to contribute to cybersickness, which we have recently discussed as a signature of the evolutionary mismatch between our sensory systems and the technological environments of the Anthropocene [5, 6]?

In this article, we argue that all of these questions can be addressed within a common computational framework provided by active inference [7, 8]. The argument has a closely related engineering counterpart in immersive technology. Michael Abrash, who has led virtual reality research at Valve and later at Oculus and Meta Reality Labs, has consistently framed the goal of immersive technology not as replication of the physical world but as satisfaction of the perceptual system: “our conscious minds never actually interact with the real world. Instead, we interact with signals from sensors”, humans are “inference machines, not objective observers”, and “virtual reality doesn’t have to replicate the real world in order to work. It just has to provide the right inputs […] to satisfy the relevant senses” [9–11]. The technical requirements associated with presence that emerged from this view (field of view, display persistence, refresh rate, tracking precision, and motion-to-photon latency of about twenty milliseconds) were identified by investigating which perceptual subsystems must be satisfied before the brain accepts the displayed world as the place one is in, a process occurring “well below conscious awareness” [11]. This is an operational description of a brain that infers the causes of its sensations under a generative model, and that accepts the displayed world as the place one is in when the hypothesis that one is in that world explains the sensory input better than the hypothesis that one is wearing a display in a room. The free energy principle and active inference provide a framework for formalizing this interpretation [7, 8, 12], and predictive processing accounts of presence have already been proposed in the context of interoception [13, 14], sensorimotor contingencies [15, 16], phenomenal transparency [17, 18], body ownership [19, 20], and the brain as a simulator [21]. Most recently, it has been argued that the sense of presence in virtual reality and the sense of reality in everyday perception are mechanistically continuous [22], and electroencephalographic prediction error signatures have been used to detect implausible interactions in virtual reality [23–26].

What is missing, is a formal treatment that (a) represents the actual and the induced reality as competing hypotheses within one generative model, (b) derives the role of attention from precision, thereby explaining the attention taxonomies of the two-competitor model rather than assuming them, (c) integrates affect, motivation, and effort through expected free energy, (d) explains breaks in immersion, sensory conflict, and cybersickness within the same variables, and (e) connects to psychophysiological measurement and to the neuroergonomic design of immersive technologies. This is the goal of the present article. We first review the constructs of immersion and presence, followed by the necessary elements of the two-competitor model and of active inference (Section 2). We then develop the active inference account of induced realities (Section 3), map it onto the two-competitor model and reproduce its two scenarios in a minimal simulation (Section 4), and discuss implications for assessment, design, closed-loop systems, and neuroergonomics (Section 5) before addressing limitations and drawing conclusions.

## 2 Background

### 2.1 Immersion, presence, and induced realities

The literature on immersion is large and terminologically heterogeneous; see Agrawal et al. [27], Skarbez et al. [28], Nilsson et al. [29] for comprehensive reviews. Different theoretical approaches define immersion and its relationship to related concepts such as presence in different ways. Slater and colleagues reserve the term immersion for the objective properties of a system, the set of sensorimotor contingencies it supports, and use presence for the subjective response [30–32]. Within presence they distinguish place illusion, the illusion of being in the place depicted, from plausibility, the illusion that the events depicted are really happening, and they emphasize that both illusions persist despite the sure knowledge that they are illusions [33, 34]. Similarly, Lombard and Ditton define presence as the perceptual illusion of nonmediation [35]. In contrast, Wirth and colleagues describe spatial presence as the outcome of a two-step process in which a spatial situation model is first constructed and then accepted as the primary egocentric reference frame [36, 37]. Finally, Agrawal and colleagues, whose definition we took as our starting point in the two-competitor model, define immersion as “a state of deep mental involvement in which the subject may experience disassociation from the awareness of the physical world due to a shift in their attentional state” [27, as quoted in 1]. Related constructs include embodiment, the sense of self-location, agency, and ownership over a virtual body [38–40], flow [41, 42], and narrative transportation [1, 43, 44]. What these definitions share is that the person is situated, by their own perceptual system, in one environment rather than another; where they differ is in what they take to be primary, the technology, the attentional state, the spatial model, or the illusion of nonmediation. Meta-analytic evidence indicates that technological immersion in Slater’s sense increases presence with a medium effect size, with tracking, field of view, and stereoscopy among the most effective factors [45]. Assessment of presence and immersion has relied on physiological responses to stressful virtual environments [46], electroencephalographic correlates of spatial presence [47, 48], and counts of reported breaks in presence [49]. In this article we keep the two terms apart as follows. Immersion denotes the graded state of the person, the degree to which the induced reality, rather than the actual one, is taken to be the reality one is in, which is Agrawal’s attentional definition made quantitative in Section 3.1. Presence denotes the subjective transparency of that state, the illusion of nonmediation of Lombard and Ditton and the place illusion of Slater, which Section 3.1 identifies with the absence of prediction error at the level of the context. Slater’s immersion, the sensorimotor contingencies a system supports, is in our terms a property of the induced reality rather than of the person, and it enters the account as the evidence that the induced reality can supply. The predictive processing accounts reviewed in Section 2.4 concern presence in this sense, and they are the natural predecessors of an account in which immersion and presence are two aspects of one inference.

#### 2.1.1 On actual, physical, and induced realities

In the two-competitor model we used the term induced reality to cover the whole range of situations in which attention is directed to a reality other than the one the person is actually in, from reading a book to a fully tracked multisensory virtual environment [1]. We continue to use the terms *induced reality* here, as well as *actual reality* rather than *physical reality* for its counterpart, because this distinction allows for cases in which physically generated cues contribute to the induced reality. A film screened with a live orchestra, a situation comedy with a laughter track or with the laughter of a physically present audience, and an extended reality installation with live-played music all integrate real-world cues into the induced reality, and such cues may deepen immersion when they are congruent with it and disrupt it when they are not [50, 51]. Whether a cue is treated as part of the induced reality or as a competing signal is therefore decided by its coherence with the other elements of the scene, not by its physical origin, and once the cues are assigned, the attentional dynamics of the two-competitor model apply. This point is easily missed. In a recent position paper, Richardson, Gilchrist, and Stanton Fraser adopt a source-based interpretation and equate the actual reality with the physical world. They then conclude, from audience laughter, shared snacks, and printed books, that input from the actual reality (i.e., the physical world) can enhance immersion, which they take to contradict the orthogonal competition of the model [52]. Under this reading, the two-competitor model indeed fails. However, the term *actual reality* was chosen in the original article in preference to the more natural *physical reality* precisely to keep the two apart, but a loose, passing gloss of the actual reality as only and entirely the physical world around us and inside us [1] has invited the source-based reading. Under the model’s own criterion, congruent laughter belongs to the comedy as much as the lines of the actors; the comfort of a printed book and the company of other viewers enter through the endogenous weights, where motivation was located from the start, and the study behind the laughter example shows the congruency dependence that this criterion predicts and a source-based reading cannot explain: laughter and screams raised reported immersion only when they matched the genre of the film [53]. Ultimately, an induced reality is in this sense always embedded in the actual reality: the display is a physical object, the reader sits in a room, and the brain continues to receive sensory information from the physical environment. Nevertheless, attention is directed toward one in preference to the other. The virtue of the terminology is that it forces a theory to explain what these situations have in common, and what distinguishes them, in a single set of variables. An active inference account meets this demand naturally, because it describes every one of these situations as inference under a generative model that receives sensory evidence of varying richness, and because it makes the assignment of a cue to a reality part of the inference itself (Section 3.5).

### 2.2 The two-competitor model

The two-competitor model represents the actual reality (ar) and the induced reality (ir) as orthogonal axes of a plane in which an attentional focus vector ***a*** = *a*_ar_ ***e***_1_ + *a*_ir_ ***e***_2_ is mapped [1]. Its angle *ϕ*_*a*_ = tan^−1^(*a*_ir_*/a*_ar_) quantifies the degree of attention directed to the induced reality, with *ϕ*_*a*_ = *π/*2 indicating that attention is directed entirely to the induced reality, and its magnitude *m*_*a*_ = ∥***a***∥_2_ quantifies the total attentional allocation. The projections are sums of exogenous and endogenous contributions,

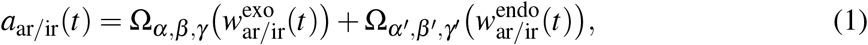

where Ω_*α,β,γ*_ (*x*) = *α/*(1 + *e*^−*βx*+*γ*^) is a sigmoidal transfer function that maps exogenous or endogenous weights into a bounded allocation of attentional resources. The model is configured to favor sufficiently strong exogenous signals, allowing abrupt and intense events to over-ride ongoing immersion. The weights themselves were defined by two parameterized maps, *w*^exo^ = Φ_***p***_(***s***(*t*)) for the stimulus-driven properties of the streams (brightness, loudness, valence, arousal) and *w*^endo^ = Ψ_***p***_(***s***(*t*)) for the goal-driven allocation that reflects internal attention, motivation, and affect, with a cognitive fatigue model [3, 4] as the worked example for Ψ_***p***_. Two scenarios illustrated the dynamics: a rewarding activity in the induced reality with a stable motivation profile, and an effortful, fatiguing activity in which motivation progressively declines, both interrupted after ten minutes by a slamming door in the actual reality that transiently pulls the attentional focus vector back toward the ar axis [1].

The model thus already contained four commitments that we now make explicit in active inference terms. First, attention is a necessary but not a sufficient condition for immersion. Second, immersion is a competition between two realities for a limited resource. Third, the competition is modulated by two modes of attentional allocation, exogenous (stimulus-driven) and endogenous (goal-directed) attention, both of which contribute to the allocation of attention between the two realities. Fourth, affect and motivation influence attentional allocation through the endogenous weights, which is where the insufficiency of attention was located. The maps Φ_***p***_ and Ψ_***p***_ were placeholders for “any model which provides this” [1]. Active inference offers a framework meeting these requirements.

### 2.3 Active inference in brief

Active inference describes perception, attention, and action as consequences of a single imperative, the minimization of variational free energy under a generative model of how sensations are caused, with expected free energy guiding the selection of future policies [7, 8, 12, 54]. In more intuitive terms, the brain continuously predicts its sensory input and compares these predictions with incoming observations. Discrepancies can be reduced either by updating beliefs about the causes of sensory input (perception) or by acting to bring sensory input into closer agreement with predictions (action) [8]. For sensory observations *o* and hidden states *s*, the generative model *p*(*o, s*) specifies the probabilistic relationships between hidden states and the observations they are expected to generate. Hidden states represent underlying conditions or causes, such as the source of a sound, which cannot be directly observed but must be inferred from sensory input. The exact posterior *p*(*s* | *o*) describes the probability distribution over hidden states given the sensory observations and is here approximated by a distribution *q*(*s*). Variational free energy measures how well beliefs about hidden states account for sensory observations. It provides an upper bound on surprise, the negative log-evidence of the observations under the model, and can be expressed as:

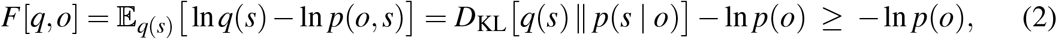

Perception reduces *F* by updating beliefs about hidden states represented by *q*(*s*), while learning adjusts the parameters of the generative model *p*(*o, s*). Action can change sensory observations by acting in the environment, thereby altering *o*. In hierarchical continuous-state formulations, the gradient of *F* includes terms involving precision-weighted prediction errors ***ζε***. Sensory prediction errors can be expressed as ***ε*** = *o* − *g*(*s*), where *g*(*s*) represents the predicted sensory input, and describe discrepancies between predicted and observed sensory input. Precision ***ζ***, the inverse of the expected variance of these errors, determines how strongly they influence inference [55–57]. Attention, in this framework, is precision optimization: increasing the precision of a sensory channel increases its influence on inference, and attentional effort may further be conceptualized in terms of the costs associated with maintaining precision estimates under uncertainty [6, 55, 58, 59].

The selection of future actions is guided by expected free energy, which evaluates possible courses of action in terms of their anticipated outcome and expected information. For a policy *ξ*, a sequence of actions, the expected free energy *G*(*ξ*) of its future outcomes decomposes as

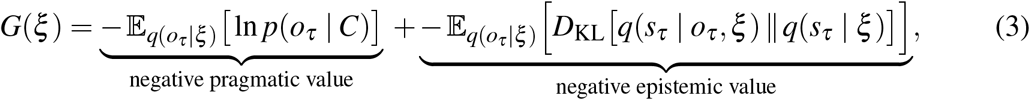

where *C* encodes prior preferences over outcomes [8, 12, 60]. In the absence of additional policy priors, policies are selected according to *q*(*ξ*) = *σ* (−*γ*_*ξ*_ *G*(*ξ*)), a softmax over policies with precision *γ*_*ξ*_; for two policies the softmax reduces to the logistic function *σ* (*x*) = 1*/*(1 + *e*^−*x*^) of the difference in expected free energy, and we use *σ* for both in what follows. The first term captures pragmatic value, reflecting how well the anticipated outcomes of a policy align with prior preferences. The second captures epistemic value, reflecting the expected information gain, or reduction of uncertainty about hidden states [58, 60, 61]. Because *G* is the sum of the negative pragmatic and the negative epistemic value, higher pragmatic or epistemic value leads to lower expected free energy. The epistemic term thereby formalizes intrinsic motivation, curiosity, and exploration, and it has been used to explain why agents seek novelty and why the epistemic value of an activity collapses once it no longer resolves uncertainty [61, 62].

Two further elements are needed. First, generative models are hierarchical and deep in time: slow, abstract states at higher levels contextualize fast, concrete states at lower levels, so that a narrative or a goal can constrain the interpretation of sensory input over seconds to hours [63– 65]. In our earlier work we represented this hierarchy as a spatiotemporal scale space with an evolutionarily entrenched, fast, and near core and a slower, reasoning-based periphery [5, 6], and we distinguished within the core a species-typical, genetically entrenched set of biases from a plastic, experience-dependent mantle [5]. Second, affect has been given a free energy account. Joffily and Coricelli proposed that valence tracks the rate of change of free energy, negative when free energy rises and positive when it falls [66]; Hesp and colleagues modeled valence as an inference based on the expected precision of one’s own action model [67]; and Kiverstein and colleagues described the feeling of grip as the experience of prediction errors being reduced at a rate that is better than expected [62, 68]. Interoceptive inference extends the framework to the body: emotions and the sense of a bodily self arise from predictions about the causes of interoceptive signals, and the precision of these predictions determines their contribution to experience [14, 69–72].

Finally, and central to what follows, Parvizi-Wayne has argued that the dichotomy between exogenous and endogenous attention becomes less clear within active inference: preferences “enslave” attention, so that a stimulus-driven capture and a goal-driven allocation are both instances of precision being assigned to the streams that matter for the expected free energy of the agent’s policies, the two differing, in our reading of his argument, in the level and time scale of the model at which the precision is set [73]. We used this argument for effortful listening in the soundscapes of the Anthropocene [6] and for sensory conflict and strain in neuroergonomics [5]. Here we use it to reinterpret the exogenous and endogenous contributions to attentional allocation in the two-competitor model.

### 2.4 Predictive processing accounts of presence

Predictive processing accounts have so far addressed presence, in the sense fixed in Section 2.1, rather than immersion. Seth, Suzuki, and Critchley proposed an early predictive processing model of presence, in which the sense of presence arises when interoceptive prediction errors are successfully suppressed by top-down interoceptive predictions. This suggests that presence depends in part on whether bodily responses can be integrated into a coherent prediction of the current situation [13]. The model was later tested in virtual reality, where only high-arousal emotional states modulated presence [74]. Building on the model, Deane et al. discussed how the sense of presence is diminished in depersonalization [75]. Seth also proposed that perceptual presence, the phenomenal sense that an object is really there, depends on the counterfactual richness of the generative model, that is, on how many sensorimotor contingencies the model encodes for the object [15, 76], an idea rooted in the sensorimotor account of perception [16] and directly relevant to Slater’s characterization of immersion as the set of valid sensorimotor contingencies a system supports [31]. Building on Metzinger’s account of phenomenal transparency, the fact that we normally experience the world rather than our model of it [77], Limanowski and Friston related this phenomenon to the precision assigned to the model itself: a model that is used for inference with high confidence is transparent, whereas a model whose precision is lowered becomes opaque, an object of attention in its own right [17, 78]. Body ownership illusions have been modeled as Bayesian causal inference over the common cause of visual, tactile, and proprioceptive signals [19, 20, 39]. Riva and colleagues described the brain as a simulator whose predictions, when matched by the technology’s predictions about the sensory consequences of the user’s actions, produce presence [21, 79], and Gonzalez-Franco and Lanier proposed a model of illusions in virtual reality based on bottom-up and top-down processing [80]. Recent electroencephalographic work has identified prediction error signatures of implausible visuo-haptic interactions in virtual reality and their modulation by presence and embodiment [23, 25, 26], complementing our own observation of a cross-modal semantic incongruency response to mismatching tactile and virtual visual experiences [24]. Singleton and Fotopoulou have recently argued that presence in virtual reality and the sense of reality in everyday perception share a predictive processing mechanism [22], and reality monitoring has been described as a comparison of subjective signal strength to a reality threshold [81, 82].

Taken together, these accounts suggest that presence can be understood as emerging from perceptual and inferential processes rather than as a separate component added to perception. To our knowledge, none of these accounts combines an explicit competition between induced and actual reality with a precision-based treatment of both attention taxonomies and an account of the affective and motivational dynamics that govern immersion. The two-competitor model contains exactly these elements in descriptive form. The following section brings the two together.

## 3 An active inference model of induced realities

### 3.1 The reality as a hidden context

We begin with the observation that the sensorium of an immersed person is never generated by a single reality. A person wearing a head-mounted display may receive visual and auditory streams from the induced reality, but also the weight of the headset, the sound of the room, the pressure of the chair, hunger, fatigue, and, crucially, proprioceptive and vestibular signals that are generated by the physical body in the physical room. A reader receives the words of the novel and, at the same time, the light of the lamp and the noise of the street. The question for the brain is therefore not which reality is present, but which reality provides the best explanation of the sensory signals it is currently sampling and the best basis for the policies it is currently pursuing.

We formalize this by introducing a hidden context variable *c* ∈ {ar, ir} as one of the slow, abstract states that, in the hierarchy of Section 2.3, contextualize the faster states and the observations below them,

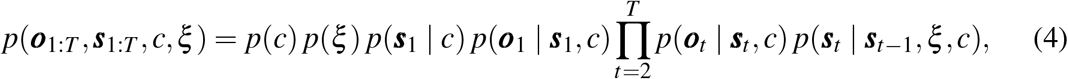

in which ***o***_1:*T*_ = (***o***_1_, …, ***o***_*T*_) and ***s***_1:*T*_ denote the sequences of observations and hidden states over the *T* time steps of an episode, and in which the context selects the likelihood mapping *p*(***o***_*t*_ | ***s***_*t*_, *c*) and the transition dynamics *p*(***s***_*t*_ | ***s***_*t*−1_, *ξ, c*), that is, the sensorimotor contingencies and the narrative and causal structure that the agent uses to predict its sensations. If the induced reality provides the relevant context (*c* = ir), predictions are based on the induced reality’s model: head rotation predicts a rotation of the displayed scene, a virtual approach predicts a change in binaural cues, and a plot predicts the next event. Under *c* = ar, the model of the actual reality is engaged: head rotation predicts corresponding shifts of the room one is in, and the plot is irrelevant. Both models make predictions about the same observation vector ***o***_*t*_, which we partition into modality-specific streams 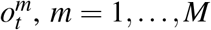, including exteroceptive streams delivered by the induced reality, exteroceptive streams that only the actual reality generates, and bodily streams (proprioception, vestibular signals, interoception) that are generated by the physical body under either context. Figure 1 summarizes the structure.

**Figure 1:**
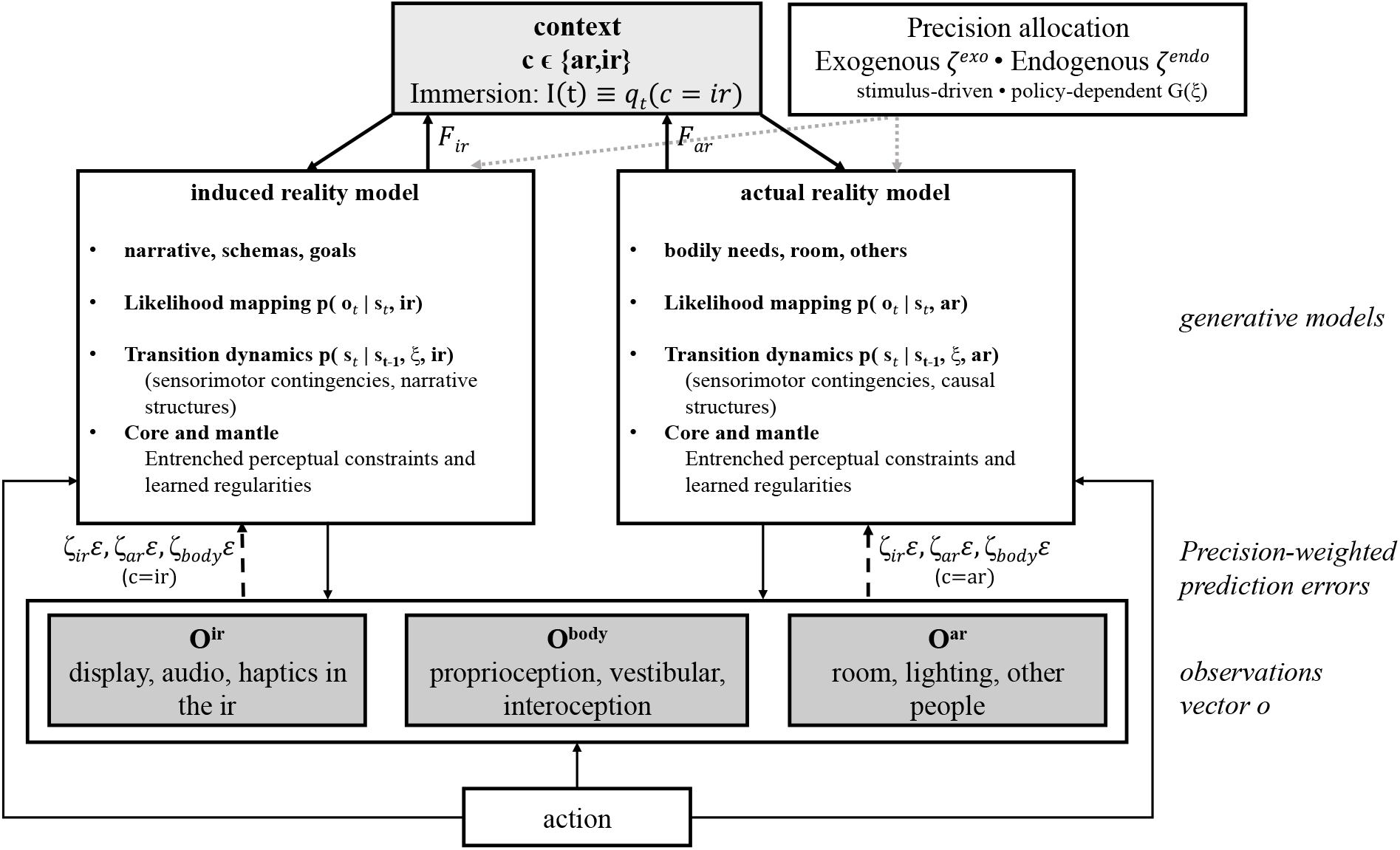
Schematic of the generative model of induced realities. A context variable *c* selects which of two models, that of the induced reality (ir) or that of the actual reality (ar), is used to predict a shared observation vector consisting of ir-specific streams, ar-specific streams, and bodily streams that are generated by the physical body under either context. Precision-weighted prediction errors contribute to the variational free energy of each context, *F*_ir_ and *F*_ar_, which informs context inference and the degree of immersion *I*(*t*) = *q*_*t*_(*c* = ir). Solid arrows denote predictions, dashed arrows precision-weighted prediction errors and dotted arrows precision control.

The two context-dependent models are not entirely independent, but share parts of their generative structure, with context-specific differences in the likelihood mappings and transition dynamics. One implication of this shared structure is that an induced world need not be constructed from scratch, an idea closely related to Ryan’s principle of minimal departure. According to this principle, fictional worlds are assumed to resemble the actual world as closely as possible, departing from it only where the narrative requires. A reader encountering a fictional world in which sorcerers exist will still assume that gravity operates as usual, even if the author never mentions it [83]. This helps explain why a novel can evoke a rich imagined world from relatively little sensory input by drawing on existing knowledge. This shared generative structure also provides a possible basis for interactions between the two realities: prior experience shapes the interpretation of induced worlds, while learning within an induced reality may update knowledge or skills that subsequently transfer to the actual reality. For example, information encountered in fiction can influence real-world beliefs [43, 84, 85], and skills acquired through VR training can transfer to real-world tasks [86, 87]. Such cross-context learning is not explicitly modeled here.

Inference on the context follows from the free energy, Eq. (2), of the two sub-models. Writing *F*_*c*_ for the variational free energy obtained when the observations are explained under context *c*, the posterior over the context is

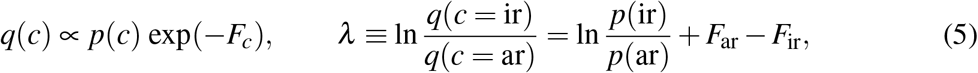

so that the log-odds *λ* of being in the induced reality are given by the difference in free energy, or equivalently in log-evidence, between the two explanations. Equation (5) is an instantaneous posterior. If the context is treated as a hidden state that changes slowly, with a transition prior *p*(*c*_*t*_ | *c*_*t*−1_) that favors staying, the prior term becomes the prediction from the previous posterior and the evolution of *λ* can be approximated by a recursive, leaky accumulation of the free energy difference over time, which is the form we use in the simulation of Section 4.2. Because free energy is a sum of precision-weighted prediction errors over streams and hierarchical levels, the difference *F*_ar_ − *F*_ir_ is dominated by the streams to which precision is assigned. We use *λ*, and its logistic transform *q*(*c* = ir) = *σ* (*λ*), as the central quantity of this article.

#### Definition of immersion

We define the degree of immersion at time *t* as the posterior probability that the induced reality is the context that best explains the current sensorium and that supports the current policy, *I*(*t*) ≡ *q*_*t*_(*c* = ir). This definition satisfies the requirements we set out for the two-competitor model. It is continuous and dynamic; it does not require introspection, since *q*(*c*) is a state of the inference machinery rather than a belief one can report on without changing it; and it covers the whole range of induced realities, because the evidence that enters Eq. (5) can come from rich exteroceptive streams, from a narrative maintained at deep levels of the model, or from both. Moreover, the framework allows the endogenous influence of affect and intrinsic motivation, central to the original two-competitor model, to be represented through preferences, expected information gain, and their effects on policy selection and attentional precision.

#### Definition of presence

Following Limanowski and Friston [17] and Metzinger [77], we define presence as the transparency of the winning model: the induced reality’s model is used to generate predictions and to guide action with high precision, while the context itself does not become an object of attention. In the terms of Eq. (5), presence corresponds to a regime in which *λ* is large and, importantly, stable, so that the induced model is used with high confidence and the context itself generates no prediction errors that would attract attention. When *λ* approaches zero, the context becomes uncertain, prediction errors at the context level grow, precision is drawn to the question “where am I?”, the mediation becomes opaque, and one notices the headset, the page, or the screen. This distinguishes presence from immersion: presence is the absence of prediction error at the context level, immersion is the outcome of the context inference. Abrash’s remark that presence “is distinct from immersion, which merely means that you feel surrounded by the image of the virtual world; presence means that you feel like you’re in the virtual world” [11] is consistent with this distinction.

#### Place illusion and plausibility

Slater’s two orthogonal illusions primarily map onto two components of *F*_ir_ [31, 33]. Place illusion is the success of the induced model at the level of sensorimotor contingencies, that is, the smallness of prediction errors in *p*(***o*** | ***s***, ir) when the agent acts: turning the head, reaching, walking. Plausibility is the success of the induced model at the level of scene and narrative states, that is, the smallness of prediction errors in *p*(***s***_*t*_ | ***s***_*t*−1_, *ξ*, ir): events unfold as the model of a coherent world predicts, and the world responds to the agent as a world would. Both contribute to the free energy difference in Eq. (5), and both can be lost independently, as Slater and colleagues have documented.

### 3.2 Precision and the two taxonomies of attention

In the two-competitor model, the exogenous and endogenous weights were given by the maps Φ_***p***_ and Ψ_***p***_. In active inference both weights are precisions, and the maps are replaced by the mechanisms that set precision. Following Parvizi-Wayne [73], we distinguish two routes of precision control, which we write as 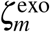 and 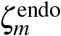 for stream *m*, not because they are different kinds of attention but because they are set at different levels of the model and on different time scales.

#### Exogenous precision

The likelihood precision of certain sensory events is high by construction of the model, and the construction is evolutionary. Abrupt onsets, looming, sudden loudness, motion in the periphery, faces, and the interruption of a regularity carry high precision because, in the environments in which the human perceptual system evolved, they were the events whose neglect was most costly [5, 6, 88–90]. These precisions are part of the genetically entrenched core of the model, they are fast, they are modality-specific, and they are largely independent of the current context. In the two-competitor model we captured this with a transfer function that gives priority to the exogenous channel, and in the fatigue scenario we represented the slamming door as a brief exogenous spike. Here the door is a high-precision prediction error in an ar-specific stream, and it enters *F*_ar_ − *F*_ir_ with a weight that the induced model cannot reduce because the induced model does not predict doors in the physical room. The sound is better explained by the actual than by the induced model. Note that the induced reality has its own exogenous events (a jump scare, a virtual explosion), and these carry the same entrenched precision, which is why they are so effective at pulling and holding attention [1].

#### Endogenous precision

The second route sets precision according to the expected free energy of policies. An agent that has selected the policy “continue reading the novel” or “continue the game” assigns high precision to the streams whose prediction errors are informative for that policy and low precision to the others. This is what Parvizi-Wayne means by preferences enslaving attention [73], it is what Feldman and Friston [55] modeled as attention, and it is the active inference reading of the biased competition and probabilistic stream selection models that we used as the basis of the two-competitor model [91–93]. Formally, we let the endogenous precision of the streams of reality *c* follow the posterior probability of the policies that act in *c*. Rather than determining endogenous precision independently for each stream, we first define a common precision at the level of each reality *c*. This context-level precision 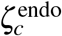 is determined by the posterior probability of policies in *c* and is assigned to the streams associated with that context. We adopt this simplification in the simulation presented, see Appendix A.

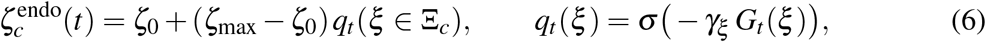

#### External and internal attention

The second taxonomy of the two-competitor model, external versus internal attention [2], does not require a third route.

Instead, it distinguishes the target of attentional selection. In external attention, precision is assigned to currently available sensory information, while in internal attention, precision is directed toward internally generated or maintained representations, such as memories, imagery, narratives, and simulations. The distinction cuts across the hierarchy of the generative model and can concern both lower sensory and higher interpretive levels. In the competition between actual and induced realities, both forms of attention can contribute to either reality. A novel, for example, engages external attention through the perception of written words and internal attention through the imagined narrative world. The relative contribution of external and internal attention to the induced reality therefore depends on the precision assigned to the respective representations.

How this precision is distributed may also depend on the sensory modality involved. Within the two-competitor model, internally generated and externally driven signals may compete for attentional precision within the same sensory modality. Vivid visual imagery could therefore reduce the attentional gain assigned to external visual input. Conversely, a sufficiently salient external stimulus might shift this balance toward the actual reality, potentially disrupting immersion more strongly when it engages the same modality as the induced representation than when it engages a different modality. The contribution of internally generated imagery may also vary between individuals, consistent with findings of reduced narrative absorption and engagement in individuals with aphantasia [94, 95].

#### Consequences

Three consequences follow. First, exogenous and endogenous precision control are not independent: exogenous precision is set by the model’s construction, endogenous precision by the model’s current policies, and the second can modulate the effect of the first only within limits set by the core. This is why one can learn to ignore the hum of an air conditioner but not a slamming door. It is the phenomenon that we have studied as long-term or attentional habituation, the drift of attention away from a repeated stimulus that is neither informative nor behaviorally relevant [96–98]. In the present terms such a stimulus carries little exogenous weight, and once its regularity has been learned it produces no prediction error and no information gain, so that the precision assigned to its stream fades and the stream drops out of the competition. In our habituation paradigm the phase locking of single-trial late auditory evoked responses declined over the repetitions of a soft tone but remained stable over the repetitions of a loud one [96–98], and we modeled this as a hippocampal comparator that detects novelty and withdraws attentional binding from familiar input [97]. A loud or aversive stimulus keeps the exogenous precision of the core and resists habituation, and a change in stimulus properties restores precision at once, which is dishabituation. Second, the competition between realities is decided by precision-weighted evidence, not by evidence alone. A headset can deliver a perfectly coherent scene and still lose the competition if the policies of the agent lie in the actual reality, and a novel can win the competition despite providing little exteroceptive evidence for the depicted world if the policies lie in the narrative. Third, the total precision that an agent can maintain is limited, and its maintenance costs effort [3, 6, 99, 100]. The magnitude *m*_*a*_ of the two-competitor model, the total attentional allocation, corresponds to the total precision currently deployed across both realities. The effort associated with maintaining and adjusting this precision allocation relates to what we have called sensory strain in the neuroergonomic context [5].

### 3.3 Affect, motivation, and the expected free energy of staying

The two-competitor model introduced affect through a fatigue module that mapped reward and effort onto the endogenous weight of the induced reality [1, 4]. Active inference offers a more general and more principled account. The endogenous precision of the induced reality follows, through Eq. (6), the expected free energy of the policies that act in it, and Eq. (3) decomposes this expected free energy into a pragmatic and an epistemic component. Both have direct readings in the phenomenology of immersion.

The pragmatic value of staying in the induced reality is the expected fulfillment of prior preferences: the reward of the game, the pleasure of the story, the social contact in a virtual meeting, the completion of a training task. The pragmatic value of leaving is the expected fulfillment of the preferences of the actual reality: eating, resting, answering the person at the door, the “increasing pressure to do something else in the actual reality” of our earlier scenario [1]. As bodily needs change during prolonged immersion, policies directed toward the actual reality may gain pragmatic value. Sustaining immersion then depends on whether the combined pragmatic and epistemic value of remaining in the induced reality continues to outweigh that of competing policies.

The epistemic value of staying is the expected information gain, that is, the reduction in uncertainty about hidden states that the agent anticipates from remaining in the induced reality. A well-designed game, a well-told story, and a well-structured training scenario keep this term high by continuously posing questions that the agent can answer through action and observation. The two-competitor scenario of a game that becomes too hard is, in these terms, a joint collapse of pragmatic and epistemic value: the reward-to-effort ratio deteriorates, and predictions are made and violated but the violations cannot be resolved by any available policy, so that expected information gain, and with it the endogenous precision of the induced reality, declines. This is a different failure from boredom, in which epistemic value collapses because nothing is left to resolve, but both tend to reduce immersion [61, 62]. The region between the two failures, in which prediction errors are continuously generated and continuously resolved at a rate that matches the agent’s capacities, is the region of flow [41, 42, 62, 68], and it is the region in which the endogenous precision of the induced reality is maximal.

Valence itself we read, following Joffily and Coricelli [66], as a function of the rate of change of free energy, positive when variational free energy falls and negative when it rises, in the spirit of the accounts that relate valence to error dynamics that are better or worse than expected [62, 67]. This gives a natural account of the affective texture of immersion. A rewarding activity in the induced reality produces a positive valence while its pragmatic value rises and prediction errors are resolved, and a neutral valence once a stable level is reached; a fatiguing activity produces a sustained negative valence as pragmatic and epistemic value decline; and a break in immersion produces a sharp negative transient, the startle, when an unexplained high-precision prediction error appears, followed by a positive transient as that error is resolved. Arousal, in turn, is the interoceptive correlate of a change in precision, which is consistent with the finding that presence is modulated by high-arousal emotional states [13, 74].

### 3.4 Breaks in immersion as Bayesian model switching

Equation (5) makes a break in immersion a Bayesian model comparison. As long as the induced model explains the precision-weighted sensorium at least as well as the actual model, *λ* remains positive and the agent stays immersed. A break occurs when a prediction error arises that the induced model cannot absorb and that carries enough precision to reverse the sign of *F*_ar_ − *F*_ir_. Three kinds of event have this property. Exogenous events in the actual reality (the slamming door) carry entrenched high precision. Violations of the induced reality’s own sensorimotor contingencies (latency, tracking loss, a hand that passes through a table, a mismatch between tactile and visual experience) generate prediction errors in streams to which the agent has assigned high endogenous precision, precisely because it is acting in the induced reality [23, 24, 26, 101]. And violations of plausibility at the narrative level (a character who behaves inconsistently, an event without cause) generate prediction errors at the deep levels of the model. The dynamics of a break follow from the precision-weighted update of *λ*. Because the rate of belief updating scales with the precision of the evidence [12, 55], a high-precision event switches the context quickly, on the time scale of an orienting response, whereas the re-accumulation of evidence for the induced reality after the event is slower, on the time scale of the endogenous precision. This asymmetry is what our earlier simulation produced with an oscillatory spike and a sigmoidal transfer function; here it is partly a property of the precision-dependent update rate and partly a property of the events themselves, since the exogenous transient is brief whereas the uncertainty it raises (what was that, and does it require action?) decays more slowly. The accumulation of evidence does not by itself make a long-immersed agent harder to break out than a newcomer; such dwell-time effects arise, in the present account, from learning, because time spent in the induced reality lets the agent learn its contingencies, which lowers *F*_ir_ and raises the disturbance required for a switch. We do not simulate learning here. Slater and Steed’s virtual presence counter, which models presence as transitions between two states driven by breaks [49], is a discrete version of this picture.

Breaks are accompanied by orienting. In the auditory modality, we have shown that the vestigial auriculomotor system, the periauricular muscles that once oriented the pinna, is activated by the direction of exogenous and endogenous auditory attention [102, 103] and by effortful listening [104], so that a non-invasive electromyogram of these muscles provides a marker of the reflexive orienting that accompanies an exogenous override. The auriculomotor system also responds to affective events [105, 106]; we return to both aspects in Section 5.1.

### 3.5 Sensorimotor coherence and causal inference

In the two-competitor model we proposed that a set of stimuli supports immersion when they “fit together in some particularly compelling way” and that this coherence is what internal attention maintains [1]. Active inference gives this a precise meaning: the stimuli fit together when they are the outputs of a single generative model in which action has predictable consequences. The induced reality’s model is engaged by the agent’s own actions. Every head movement is a test of the hypothesis *c* = ir, and the hypothesis survives only if the displayed scene responds as a world would. This is why tracking and latency dominate the technical requirements for presence [10, 11, 101, 107], why interactivity increases immersion [1, 45], and why Slater’s definition of immersion as valid sensorimotor contingencies is the right definition of what a system must provide [31, 34]. Seth’s counterfactual richness [15] adds that the sense of reality of an object increases with the number of sensorimotor contingencies the model encodes for it: an object one can walk around and pick up is more present than one that can only be looked at.

The same logic settles how a stream is assigned to a reality (Section 2). Every stream enters both *F*_ir_ and *F*_ar_ in Eq. (5), and it counts as evidence for whichever model predicts it under the current precisions, irrespective of its physical origin. The live orchestra in the concert hall is predicted by the induced model of the film and adds evidence for *c* = ir; a telephone ringing in the same hall is not, and adds evidence for *c* = ar. Congruency, in this reading, is predictability under the induced model, and the assignment is a causal inference over the source of a signal of the same kind that decides whether multisensory signals share a common cause before they are integrated [108, 109]. Multisensory integration and immersion are therefore the same computation at two levels of the hierarchy: within a reality, the brain decides whether two cues share a cause and, if they do, fuses them weighted by their precisions [108, 110]; across realities, Eq. (5) decides whether the streams share a reality. The factors that break binding at the lower level, temporal asynchrony beyond the binding window, spatial offset, and semantic incongruency, are for this reason the factors that break immersion at the upper level, which answers the question we left open in the multimodal section of the two-competitor model [1]. In this account, the assignment of sensory streams to the two contexts is itself inferred rather than given by the stimuli, and the axes of the two-competitor plane are hypotheses about the causes of the sensorium.

The bodily streams occupy a special place. Proprioception, vestibular signals, and interoception are generated by the physical body under either context, so that the induced model must explain them or must at least not be contradicted by them. A seated user whose avatar sits is consistent; a seated user whose avatar flies generates a vestibular prediction error under the induced model that cannot be resolved by action. This is the sensory conflict of cybersickness [111–113], which we have argued arises from a deeply entrenched core of the model that predicts the covariation of visual and vestibular signals and that cannot be retrained on the time scale of a session [5, 6]. Within the present model, cybersickness is free energy that neither perception nor action can reduce: the induced model predicts the visual stream but not the vestibular stream, the actual model predicts the vestibular stream but not the visual stream, and no policy in either reality resolves the conflict. Formal treatments of visuo-proprioceptive conflict [114] and of sensory attenuation and illusions [115] in active inference, and of the difference between virtual and physical head pose [116] are consistent with this reading, as is the empirical finding that presence and cybersickness are negatively related [113]: the same prediction error that causes sickness reduces *λ*.

Embodiment follows the same logic. Ownership of a virtual body arises when the commoncause hypothesis, that the seen body and the felt body are one, has more evidence than the separate-cause hypothesis [19, 39, 40, 117], and full-body illusions in virtual reality show that this inference tolerates differences in the size, age, and appearance of the body as long as a first-person perspective, spatial coincidence, and sensorimotor and multisensory contingencies are preserved [118–122]. An embodied avatar increases immersion because it moves the bodily streams from the “not contradicted” category into the “explained” category of the induced model, and it increases sensitivity to violations of sensorimotor contingencies for the same reason [26].

### 3.6 Interoception, the core, and the mantle

Two further elements of our earlier work map onto the model. First, the interoceptive predictive coding model of presence [13] concerns the bodily streams in Figure 1: presence is supported when the induced model successfully predicts, or at least does not contradict, the current interoceptive state, so that arousal is attributed to the induced world. A horror scene that raises the heart rate is consistent with the induced model, which predicts fear; a horror scene watched while worrying about an email is not, because the arousal is attributed to the actual reality.

Second, the distinction between the core and the mantle of the perceptual system [5] provides a way of considering how readily different perceptual processes can be modified through experience. The core refers to the evolutionarily shaped, species-typical architecture that constrains how sensory information is processed and weighted and, consequently, how perception and attention are directed. It includes, for example, the omnidirectional monitoring of the auditory system, vestibular–proprioceptive coupling, the priority of foveal vision, the multisensory mapping of peripersonal space, and rapid cross-modal orienting to salient events. The mantle, in contrast, comprises those aspects of perception that are more readily shaped by experience, such as learned contingencies of a particular interface, tolerance for a certain latency, or habituation to a recurring background sound [5]. The temporal window within which multisensory signals are bound is a mantle property as well. The brain maintains the perception of synchrony across modalities whose transduction latencies differ, and it expects a larger gap between the visual and the auditory component of a distant event than of a near one, as between lightning and thunder [123]; the window also narrows with training [124]. The coupling of visual and vestibular signals, by contrast, belongs to the core. For a theory of immersion, this distinction is relevant in both the induced and the actual reality, because the extent to which a stimulus can be ignored, accommodated, or given attentional priority determines whether it supports the induced reality or competes with it. Some components of attention show substantial biological constraints and inter-individual genetic variation; for example, genetic variants have been associated with differences in visual attention [125], and genetic influences have also been reported for the emotion-modulated startle response [126]. This does not imply that such processes are unchangeable, but rather that not all perceptual and attentional tendencies are equally susceptible to adaptation. For immersive design, this means that not every discrepancy should be expected to become equally tolerable with experience. A VR game can assume that players will learn an initially unfamiliar mapping between controller input and virtual action, but it should not assume that repeated, unexpected sounds from the actual environment will become equally easy to ignore. Conversely, sudden onsets can deliberately be used within the induced reality to direct attention toward relevant events, while comparable events originating from the actual environment may compete with the induced reality and potentially disrupt immersion. An immersion system should therefore accommodate relatively constrained perceptual tendencies while exploiting the adaptability of more experience-dependent processes. This asymmetry forms the basis of the design principle discussed in Section 5.2.

## 4 Reading the two-competitor model through active inference

### 4.1 Correspondence

Table 1 lists the elements of the two-competitor model and their counterparts in the present account. The correspondence is not one of replacement: the two-competitor model remains the appropriate level of description for experimental design and for the graphical intuition, and its quantities can be computed from the active inference variables. The endogenous weights 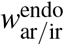 are the endogenous precisions of Eq. (6), set by expected free energy; the exogenous weights 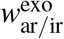 are the entrenched likelihood precisions of the events currently present in each reality; the transfer functions Ω implement a sigmoidal saturation of precision into a bounded resource allocation; the projections *a*_ar*/*ir_ and hence *ϕ*_*a*_ and *m*_*a*_ follow from Eq. (1). What the active inference account adds is the quantity *q*(*c* = ir), the perceptual outcome of the competition, which is related to the attentional focus vector but not identical to it: the angle *ϕ*_*a*_ describes where precision is currently deployed, *q*(*c* = ir) describes which reality has won the explanation. The two can dissociate, for instance immediately after a break, when precision has returned to the induced reality but the context has not yet been restored.

**Table 1.**
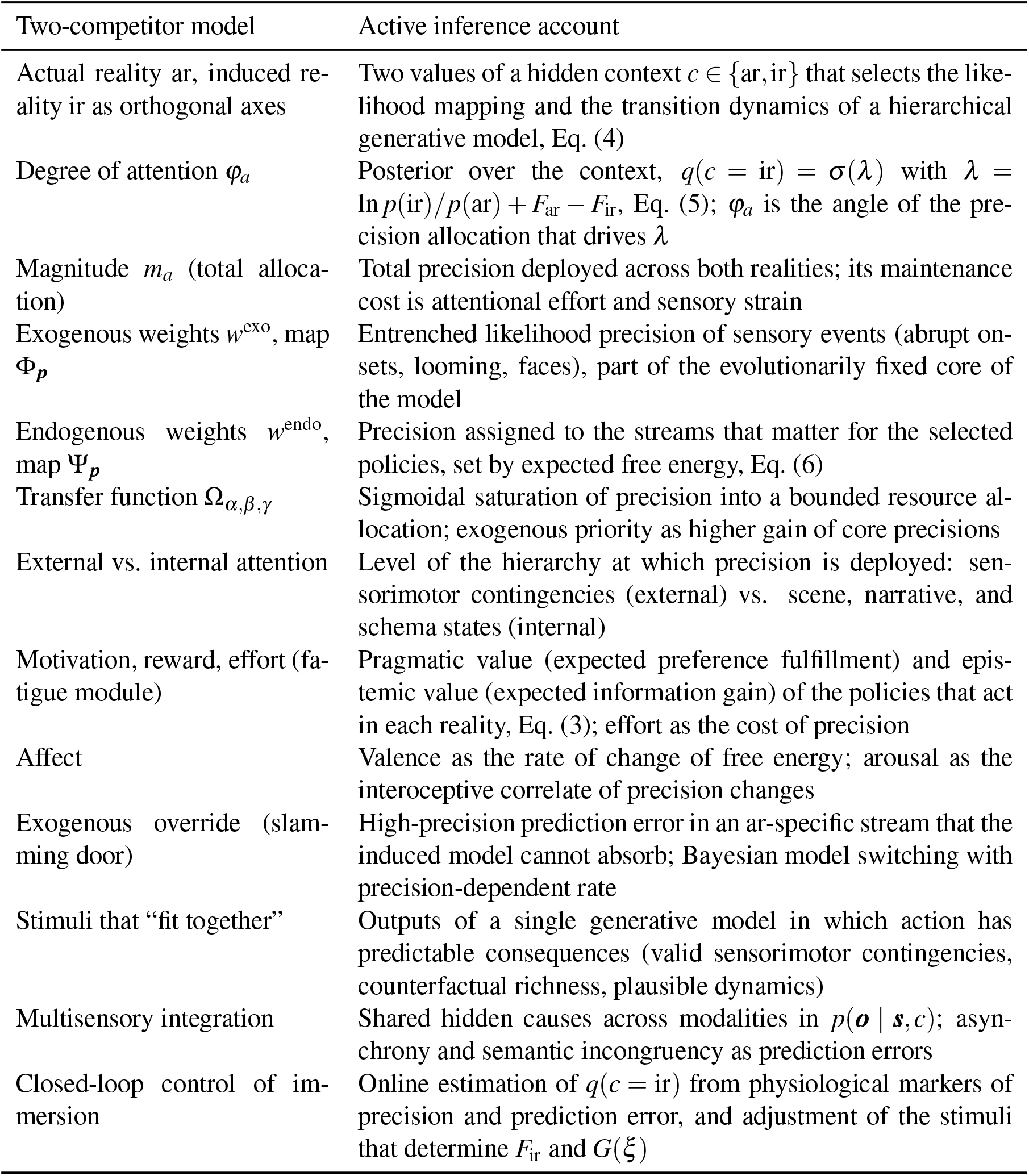
Correspondence between the two-competitor model [1] and the active inference account of induced realities.

| Two-competitor model | Active inference account |
|---|---|
| Actual reality ar, induced re-ality ir as orthogonal axes | Two values of a hidden context *c* ∈ {ar,ir} that selects the like-lihood mapping and the transition dynamics of a hierarchical generative model, Eq. (4) |
| Degree of attention *φ<sub>a</sub>* | Posterior over the context, *q*(*c* = ir) = *σ*(*λ*) with *λ* = ln *p*(ir)/*p*(ar) + *F*<sub>ar</sub> − *F*<sub>ir</sub>, Eq. (5); *φ<sub>a</sub>* is the angle of the pre-cision allocation that drives *λ* |
| Magnitude *m<sub>a</sub>* (total alloca-tion) | Total precision deployed across both realities; its maintenance cost is attentional effort and sensory strain |
| Exogenous weights *w*<sup>exo</sup>, map Φ<sub>***p***</sub> | Entrenched likelihood precision of sensory events (abrupt on-sets, looming, faces), part of the evolutionarily fixed core of the model |
| Endogenous weights *w*<sup>endo</sup>, map Ψ<sub>***p***</sub> | Precision assigned to the streams that matter for the selected policies, set by expected free energy, Eq. (6) |
| Transfer function Ω<sub>*α*,*β*,*γ*</sub> | Sigmoidal saturation of precision into a bounded resource al-location; exogenous priority as higher gain of core precisions |
| External vs. internal attention | Level of the hierarchy at which precision is deployed: sen-sorimotor contingencies (external) vs. scene, narrative, and schema states (internal) |
| Motivation, reward, effort (fa-tigue module) | Pragmatic value (expected preference fulfillment) and epis-temic value (expected information gain) of the policies that act in each reality, Eq. (3); effort as the cost of precision |
| Affect | Valence as the rate of change of free energy; arousal as the interoceptive correlate of precision changes |
| Exogenous override (slam-ming door) | High-precision prediction error in an ar-specific stream that the induced model cannot absorb; Bayesian model switching with precision-dependent rate |
| Stimuli that “fit together” | Outputs of a single generative model in which action has predictable consequences (valid sensorimotor contingencies, counterfactual richness, plausible dynamics) |
| Multisensory integration | Shared hidden causes across modalities in *p*(***o*** \| ***s***, *c*); asyn-chrony and semantic incongruency as prediction errors |
| Closed-loop control of im-mersion | Online estimation of *q*(*c* = ir) from physiological markers of precision and prediction error, and adjustment of the stimuli that determine *F*<sub>ir</sub> and *G*(*ξ*) |

### 4.2 An illustrative simulation

To illustrate how the active inference account can reproduce the behavior of the two-competitor model and to make the dependencies explicit, we implemented a simplified version of the model and repeated the two scenarios of Strauss et al. [1]: a rewarding activity in the induced reality and an effortful, fatiguing activity, both interrupted after ten minutes by a slamming door in the actual reality. The implementation follows Eqs. (3) to (6) with the following simplifications. Rather than computing pragmatic and epistemic values from a full generative model, we specified how these values change over time for the policies associated with each reality: in scenario A the pragmatic value of the induced reality rises to a stable level and its epistemic value remains constant, whereas in scenario B both decline as the game becomes too hard, and in both scenarios the pragmatic value of the actual reality rises slowly with bodily needs. Endogenous precisions followed the policy posterior according to Eq. (6). Exogenous precisions were modeled, as in the original, by oscillatory functions, with an up-chirp in the induced reality of scenario B to represent the increasingly rapid stimulation of the difficult game, and the door was represented as a transient increase in the exogenous precision of an ar-specific stream together with a transient epistemic value of the actual reality (the question “what was that?”). Rather than computing the context posterior from a full generative model at each time step, we used a simplified dynamic update of the context log-odds *λ*. The context log-odds express the relative probability of the induced versus the actual reality on a logarithmic scale, and were updated by accumulating the difference in precision-weighted sensory evidence between the two realities. This accumulation was leaky, meaning that earlier evidence gradually lost its influence, and included a prior term informed by the posterior probabilities of the current policies. The update rate was proportional to the total precision, so that more strongly weighted evidence could lead to faster changes in the inferred context. The two-competitor angle *ϕ*_*a*_ was recovered from the same precisions through the original transfer functions and parameters of Strauss et al. [1]. Valence was estimated from the smoothed negative rate of change of a simplified free energy measure. This proxy combined the expected free energy of the policy mixture with a term representing unexpected exogenous events in the reality that is currently considered less likely. The update equations and parameters are given in Appendix A. The code that generates Figures 2 and 3 can be downloaded together with the Induced Reality Simulator, a browser application in which all parameters and events of the model can be varied and the resulting time courses inspected directly; the link is given in the statement on data and code availability at the end of the article.

**Figure 2:**
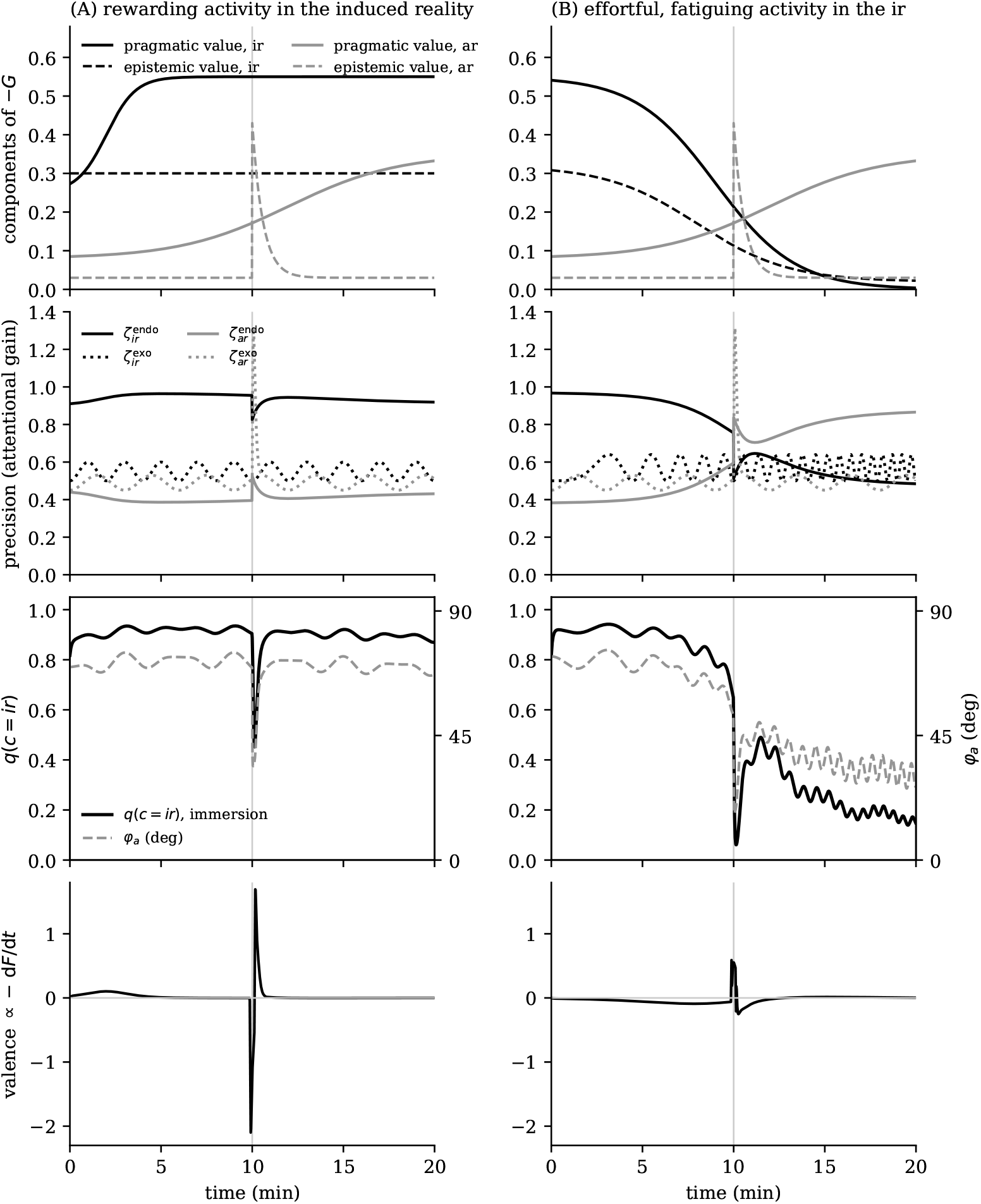
Active inference reproduction of the two scenarios of the two-competitor model. Left column: a rewarding activity in the induced reality (ir). Right column: an effortful, fatiguing activity. A slamming door in the actual reality (ar) occurs at *t* = 10 min (vertical line). First row: pragmatic and epistemic value of the policies that act in each reality, that is, the components of the negative expected free energy −*G*. Second row: endogenous precisions, which follow the policy posterior, and exogenous precisions, which follow the entrenched likelihood precision of sensory events; the door is a transient of 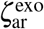. Third row: the degree of immersion *q*(*c* = ir) (solid) and the two-competitor angle *ϕ*_*a*_ recovered through the original transfer functions (dashed). Fourth row: valence as the negative rate of change of the free energy proxy. In scenario A, immersion is high and stable, the door produces a fast break and a slower recovery, and valence is mildly positive while the pragmatic value of the induced reality rises and neutral thereafter, apart from the startle transient at the door. In scenario B, the collapse of pragmatic and epistemic value withdraws endogenous precision from the induced reality, immersion declines before the door, the break is deeper and the recovery incomplete, and valence is negative until the induced reality is abandoned. Compare with Figure 3 of Strauss et al. [1].

**Figure 3:**
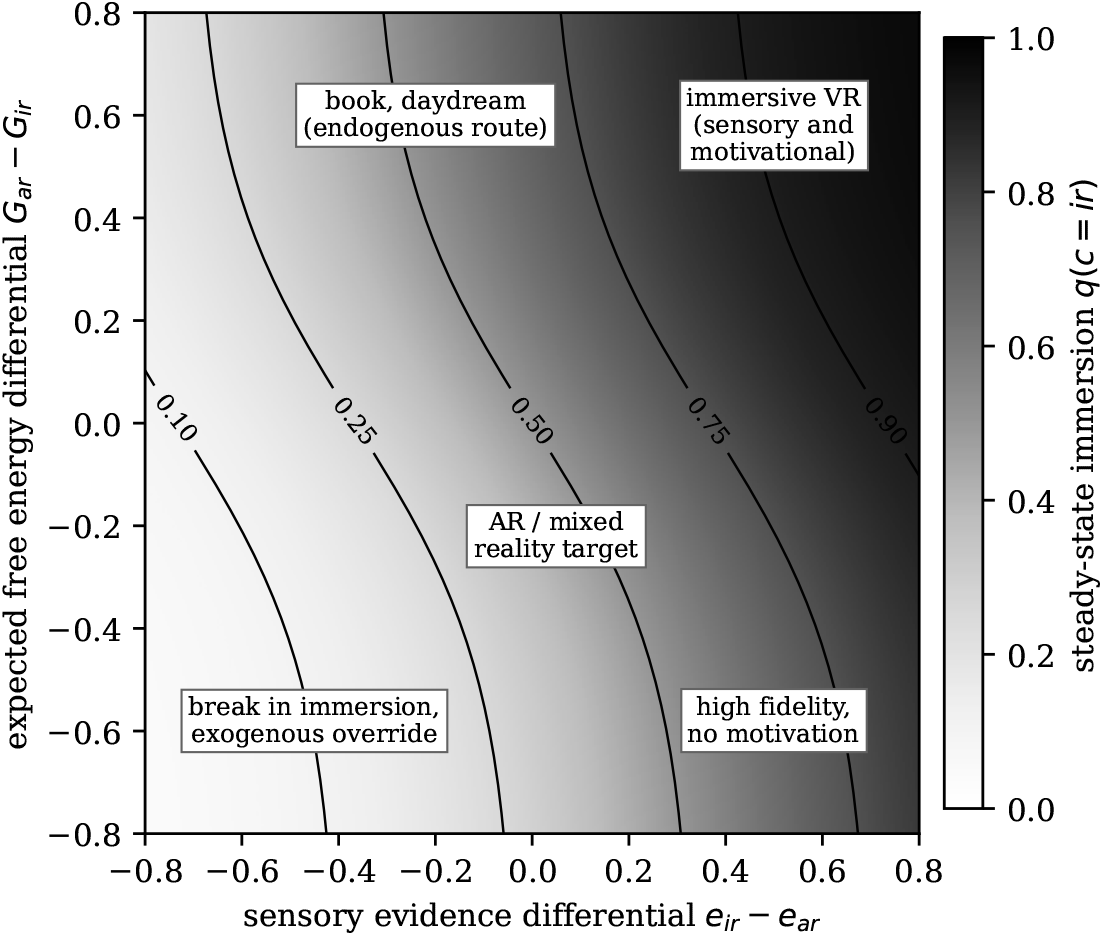
Steady-state degree of immersion *q*(*c* = ir) as a function of the difference in precision-weighted sensory evidence between the two realities (horizontal axis) and the difference in expected free energy between the policies that act in the actual and in the induced reality (vertical axis), for the parameters of the simulation. The two differentials are treated as independent arguments of the steady-state log-odds, although in the full model the endogenous component of the sensory evidence itself depends on the policy posterior. The map organizes the situations covered by the term induced reality: immersive virtual reality combines sensory and motivational evidence; a book or a daydream relies on the endogenous route with weak or negative sensory evidence; a high-fidelity system without motivation and a break in immersion occupy the lower half; and augmented and mixed reality target an intermediate, stable superposition rather than a win.

Figure 2 shows the result. The account reproduces the qualitative behavior of the original simulation: a stable, high degree of immersion in the rewarding scenario with a transient break at the door, and a progressive loss of immersion in the fatiguing scenario with a deeper break and an incomplete recovery. The two-competitor angle *ϕ*_*a*_, computed from the same precisions, follows *q*(*c* = ir) closely but not identically: *ϕ*_*a*_ is an instantaneous function of the precisions and therefore responds without delay to the exogenous transient, whereas *q*(*c* = ir) must reaccumulate evidence after the transient has passed, so that the two quantities dissociate briefly around every break. The simulation also makes explicit what the original model left implicit: in scenario B the loss of immersion is caused by the collapse of pragmatic and epistemic value, which withdraws endogenous precision from the induced reality before any exogenous event occurs, and the door only accelerates a switch that the expected free energy had already prepared.

Figure 3 shows the steady-state degree of immersion as a function of the two differentials that enter *λ*, the difference in precision-weighted sensory evidence and the difference in expected free energy between the two realities. The map organizes the situations that the term induced reality covers. Immersive virtual reality with a motivating task occupies the upper right, where sensory and motivational evidence agree. A novel or a daydream occupies the upper left, where the sensory evidence favors the actual reality but the policies lie in the induced one; immersion here is real but bounded, and it depends on the endogenous precision of the actual reality’s streams being held at its floor, which is why reading is so easily disrupted. A high-fidelity system without a motivating task occupies the lower right and produces only moderate immersion, a situation familiar to anyone who has demonstrated a headset to an uninterested visitor. A break in immersion occupies the lower left. Augmented and mixed reality, finally, target the center of the map: their purpose is a stable superposition of the two realities rather than a win of either, so that the design goal is not *ϕ*_*a*_ → *π/*2 but a stable intermediate *q*(*c*) in which the augmenting content is explained as part of the actual reality. The map is also the quantitative form of the hypothesis that attention is necessary but not sufficient for immersion [1]: precision determines which streams count as evidence, but only the expected free energy of policies that lie in the induced reality turns attended evidence into belief, and the lower right of the map shows how little sensory evidence achieves on its own. We return to this point in Section 5.2.

## 5 Implications

### 5.1 Assessment of immersion

Subjective assessments of immersion and presence commonly rely on self-report questionnaires administered after the immersive experience, such as the Presence Questionnaire (PQ), the Immersive Experience Questionnaire for Film and TV (Film IEQ) or the Immersive Experience Questionnaire (IEQ) [127–129]. While these measures are useful for capturing participants’ conscious experience, they are mostly experience-specific and inherently retrospective and may therefore be influenced by recall biases, demand characteristics, and individual differences in the interpretation of questionnaire items. Moreover, post-hoc questionnaires provide limited temporal resolution and may fail to capture brief fluctuations or breaks in immersion, whereas continuous or in-situ self-reports can themselves interfere with the immersive experience [27]. In the two-competitor model we argued that the objective assessment of immersion should combine measures of the attentional focus vector’s magnitude with measures of its direction, and that the latter requires the co-registration of events in both realities with evoked and event-related electrophysiological responses [1]. The active inference account sharpens this program by linking physiological measures to the precision allocation, prediction errors and the competition between actual and induced realities.

Event-related responses to violations of regularity, the mismatch negativity and the P3a [130, 131], are established correlates of precision-weighted prediction error [55, 56]. Probing both realities with matched irregular events (a deviant tone in the room, a deviant event in the virtual scene) and comparing the event-related responses may provide an indirect index of relative precision allocation, provided that differences in prediction error and sensory processing are appropriately controlled. The N1 attention effect and other sensory gain effects [132] may provide complementary indices of relative precision allocation at the level of early sensory processing. Our cross-modal semantic incongruency response to mismatching tactile and virtual visual experiences [24] and the prediction error signatures of implausible visuo-haptic interactions reported by Gehrke, Gramann, and colleagues [23, 25, 26] index sensory prediction errors under the induced model, which contribute to the term *F*_ir_ in Eq. (5).

Alpha power, which we and others have associated with internal attention and with the gating of sensory input [1, 48], indexes the attenuation of precision assigned to external sensory streams and is therefore particularly relevant for assessing internally maintained immersion, such as narrative absorption. The electromyogram of the vestigial auriculomotor system deserves a separate comment, because it gives access to both components that the account requires for immersion. The extrinsic periauricular muscles indicate the direction of exogenous and endogenous auditory attention [102, 103], raise their tonic activity during sustained selective listening [133], index listening effort [104], and show a brief suppression of the superior auricular muscle during reflexive orienting [134], so that their activity tracks the allocation of precision between the two realities and the cost of maintaining it. The intrinsic muscles tragicus and antitragicus, in turn, increase their activity to positively and to negatively arousing audiovisual events and correlate with subjective arousal [106], and the postauricular reflex to a probe sound is larger after a fearful than after a neutral face in far peripheral vision [105]. The same muscle system therefore also reads the affective side of the response, arousal, which the account relates to changes in precision (Section 3.3); the valence-related activity of the corrugator and zygomaticus muscles [135] adds the second affective dimension, which the account reads as the rate of change of free energy. We have argued elsewhere that peripheral channels of this kind resolve affective processing at the timescale of the events that evoke it, together with its moment-to-moment coupling to attention and effort, which neither an observer nor self-report can resolve without interrupting the state that is to be measured [136]. Immersion, which requires attention and affective resonance together, is a case in point. The same work also marks the limits of such channels. Auricular activity is motor activity that is shaped by stimulus features, attention, and effort as well as by arousal, so that a rise is compatible with a shift of attention, with effort, and with excitement, and neither autonomic measures nor facial movements carry a fingerprint of emotion categories [137, 138]. In the present account this ambiguity is the expected form of the data rather than a defect, because attention, effort, and arousal are consequences of the same precision dynamics, and the markers are to be read as evidence in the estimation problem of Section 5.3, as a dimensional state with its uncertainty, rather than as labels. Together with skin conductance and heart rate [46, 139], the auricular electromyogram might then provide an estimate of the interoceptive precision changes that accompany a break in immersion, and it might separate a break that is only an orienting response from one that is also an affective event. The virtual presence counter of Slater and Steed [49] can be interpreted as a count of model switches, while the recovery time after each break, which the active inference account predicts to depend on endogenous precision, provides an additional dynamic indicator of the depth of immersion. Beyond the number of breaks, their temporal dynamics may provide further insight into the processes underlying immersion: while a high-precision prediction error can rapidly disrupt immersion, recovery requires the re-accumulation of evidence for the induced reality, a process that depends in part on endogenous precision allocation and might therefore serve as a marker.

Immersion is inherently individual and should be considered as such: even a realistic and internally coherent virtual environment does not necessarily produce the same degree of immersion across individuals. From an active inference perspective, Parvizi-Wayne [73] argues that attention is shaped by expectations and preferences that extend beyond an immediate goal. Accordingly, the allocation of attention toward an induced reality may depend on personal experiences, long-term preferences, and emotional relevance. Bouizegarene et al. [140] describe narratives as integrative structures at higher levels of generative models, organizing predictions about temporally and causally related events. These predictions draw on individual experiences and culturally shared expectations, while new events can in turn modify existing beliefs. Consequently, the same story may be interpreted differently depending on the expectations and beliefs a person brings to it. Experimental evidence for this comes from several studies examining differences in neural responses to narratives. Presenting the same ambiguous story with different interpretive contexts beforehand led to divergent fMRI responses between groups, while responses were more similar among individuals who shared the same interpretation [141]. Importantly, such differences are not limited to externally provided interpretive cues. Finn et al. [142] found that individual differences in trait paranoia were associated with distinct neural responses to the same narrative. More recent findings indicate that belief and disbelief influence neural synchronization during narrative processing [143], while greater similarity in narrative understanding is associated with more similar neural responses [144]. These differences in prior knowledge and interpretation may also influence the degree of immersion. For example, prior knowledge or experience relevant to a story’s themes has been associated with greater narrative transportation [44]. Individuals also differ in their general tendency toward absorption, a disposition assessed by the Tellegen Absorption Scale and associated with hypnotic susceptibility and aspects of openness to experience [145].

Three experimental paradigms follow. First, a graded violation paradigm, in which the sensorimotor contingencies of the induced reality are parametrically degraded (latency, tracking noise, visuo-haptic offset) while prediction error markers and the reported break rate are recorded, tests the prediction that increasing violations of sensorimotor contingencies are accompanied by stronger physiological prediction-error responses and an increased rate of breaks in presence, and that the tolerance is larger for mantle contingencies than for core contingencies. Second, a dual-probe paradigm, in which matched exogenous probes are delivered in both realities at unpredictable times, tests the prediction that the ratio of the evoked responses provides an index of *ϕ*_*a*_ and that the auriculomotor response to an ar probe is larger when the user is more immersed, because the probe is then more surprising under the current model. Because the postauricular reflex is also modulated by the affective content of the preceding event [105], the probes must be affectively neutral and matched across the two realities, and the tonic activity of the intrinsic muscles provides a covariate for arousal from the same recording. Third, an epistemic value paradigm, in which the difficulty or the novelty of the task in the induced reality is manipulated to produce boredom, flow, and overload, tests the prediction that immersion follows the expected information gain and that effort markers [3, 104] dissociate from immersion markers in the overload condition, where effort is high and immersion declines.

### 5.2 Design principles for immersive technology

Abrash’s dictum that a virtual reality only has to provide the right inputs to satisfy the relevant senses [9, 10] becomes, in the present account, a hierarchy of requirements ordered by precision. The requirements that Abrash and colleagues derived for presence a decade ago [10, 11], a wide field of view, low persistence, high refresh rate, precise tracking, and low motion-to-photon latency, all concern the fast, entrenched core of the perceptual system: the sensorimotor contingencies between self-motion and optic flow, the vestibular-visual coupling, and the temporal precision of the visual system. These contingencies carry high, non-negotiable precision, so that their violation produces prediction errors that no narrative can absorb. Requirements that concern the mantle, such as the fidelity of textures, the realism of avatars, or the naturalness of a particular interaction metaphor, are negotiable and can be traded against cost, because the corresponding precisions are learned and can be lowered by experience. The recent research directions of Reality Labs that Abrash described, varifocal displays that satisfy the accommodation-vergence contingency, near-retinal resolution, wide field of view, and electromyographic wristbands for low-friction action [9], are all attempts to satisfy core contingencies, and the account explains why they matter more than photorealism.

The account also gives a reason for two design intuitions that are widely shared but rarely justified. First, multisensory congruency matters more than unisensory fidelity [1, 146, 147]: congruent streams are explained by a shared hidden cause and therefore reduce *F*_ir_ jointly, whereas an incongruent stream, however good in isolation, adds a prediction error. Second, narrative coherence matters even in sensory-driven immersion, because the deep levels of the induced model are the ones that maintain the endogenous precision of its streams. A world that responds to the user, in which events have causes and in which the user’s actions have consequences, keeps the epistemic value of the policies that act in it high. Adaptive difficulty, which game designers have long used to keep players in flow, is in these terms the regulation of epistemic value, and it can be made explicit by estimating the rate at which the user resolves uncertainty and adjusting the task accordingly.

Augmented and mixed reality present a different design goal, which the map of Figure 3 makes explicit. Here the augmenting content must be explained as part of the actual reality: it must respect the sensorimotor contingencies of the room, be occluded by real objects, cast shadows, and, above all, not compete with the actual reality’s streams for precision when the actual reality matters, as it does when walking through traffic or operating a machine. The design goal is a stable, intermediate *q*(*c*), a superposition in which the augmenting content inherits the transparency of the actual reality rather than competing with it. Abrash’s description of contextual artificial intelligence in augmented reality glasses, which infers what the user needs from the situation and delivers it with minimal friction [9], is a description of a system that manages the precision of its own outputs so as to reduce, rather than to increase, the free energy of the user’s model of the actual reality. This connects to our neuroergonomic proposals for attention-aware modality selection in digital operating rooms and industrial settings [5, 148**?**]: the system should route critical information to the modality and the location in the scale space where precision is currently available, and it should refrain from adding streams that would have to be suppressed.

### 5.3 Closed-loop control of immersion

In the two-competitor model we proposed closed-loop systems that use physiological measures to maintain immersion at an intended level, for instance by sending an appropriate stimulus when a drift of attention away from the induced reality is detected [1]. The active inference account turns this proposal into an estimation and control problem. The state to be estimated is *q*(*c* = ir), together with the precisions and the expected free energy terms that drive it; the observations are the physiological markers of Section 5.1; and the control variables are the stimulus properties that determine *F*_ir_ (coherence, latency, congruency), the events that determine exogenous precision (salient events in the induced reality), and the task properties that determine pragmatic and epistemic value (reward, difficulty, novelty). A controller that keeps *λ* in a target range by acting on these variables is a neuroadaptive system in the sense of our work on empathetic, context-sensitive human-technology interaction [5, 148]. We have recently specified such a loop for affect-aware systems in general: a generative model of affect grounded in active inference carries a state estimate across time, predicts the effect of the system’s own actions on that state, and selects actions by expected free energy, under bounded loop gain and with human oversight [136]. An immersion controller is a special case in which the estimated state includes *q*(*c* = ir) and the objective is a target level of immersion, and it inherits the conditions under which such a loop is steerable rather than manipulative, in particular an objective set by the user and explicit, revocable consent for every sensing channel. The auricular channel of Section 5.1 is attractive for this purpose because electrodes in and around the ear, which can be integrated into headsets and earbuds, deliver the direction of attention, effort, and arousal from one site [103, 136]. The estimation side of the loop is what we have proposed, for neuroergonomic design in general, as a surrogate human digital twin: a generative model of the latent processes behind embodied perception and cognition that updates its beliefs from physiological and behavioral channels weighted by their reliability, and that permits counterfactual tests in which priors or inputs are changed [149]. With the model of Section 3 as its computational backbone, such a twin estimates *q*(*c* = ir) together with the precisions and the expected free energy terms that drive it, and it allows a design variant, an added latency, an incongruent stream, a change in task difficulty, to be tested on the twin before it is tested on people. Two features of the account are relevant for such a controller. First, the asymmetry between the fast break and the slow recovery implies that prevention is more effective than repair: a controller should reduce the probability of high-precision ar events (by noise management and by scheduling) rather than react to them. Second, the collapse of epistemic value in the overload and the boredom regimes implies that the controller should monitor the rate of uncertainty resolution, for which effort markers and performance are the natural observations, and adjust difficulty before immersion declines. For therapeutic applications, in which the level of immersion or of arousal is itself the therapeutic variable, as in exposure therapy or in rehabilitation [1, 21, 150], the same controller can be used to hold *q*(*c* = ir) or the interoceptive precision at a prescribed level.

### 5.4 Neuroergonomics and the Anthropocene

We have argued that the technological environments of the Anthropocene impose sensory conflict and sensory strain on a perceptual system that evolved for different environments [5, 6]. Immersive technologies are the most extreme instance of such environments, because they replace, rather than merely alter, the sensory statistics to which the core is tuned. The present account locates the two problems in the same variables. Sensory conflict is free energy at the core that neither perception nor action can reduce; it is what makes an induced reality sickening. Sensory strain is the sustained cost of maintaining the precision allocation that keeps *λ* positive against a competing reality; it is what makes an induced reality tiring. Both are increased by every stream that the induced model must suppress rather than explain, and both are reduced by design that respects the core. The account therefore supports the neuroergonomic principle we have proposed, that technology design should start from the sensory statistics to which the core is tuned [5], and it adds a prediction: an induced reality that explains the bodily streams (through embodiment, through consistent self-motion, through interoceptive plausibility) is more immersive and less straining than one that merely suppresses them.

### 5.5 Reality monitoring

Presence, in the definition of Section 3.1, is the absence of prediction error at the context level. Slater et al. noted that a perfect virtual reality, indistinguishable from the physical world, would abolish presence as a distinct experience because the participant would not know that a virtual reality was in use [34]. In the present terms, a perfect induced reality is one in which *F*_ir_ ≤ *F*_ar_, so that no evidence remains that could restore *q*(*c* = ar). Reality monitoring, the capacity to distinguish perception from imagination and mediated from unmediated experience, depends on such evidence [22, 81, 82]. One design consequence follows: a route back to the actual reality remains available at low cost only if there is a class of events in the actual reality whose precision remains sufficiently high during immersion.

## 6 Limitations and outlook

The account presented here is a theoretical framework, and several of its elements are simplifications that future work must address. First, we have treated the context as a binary variable. The examples of Section 2.1 do not require more, because a cue of any physical origin is assigned to one of the two realities by its coherence with the scene. Real situations do, however, involve nested realities (a virtual meeting inside a physical office, a story within a game, a memory within a daydream) in which more than two explanations compete and the context is itself hierarchical. The extension to a set of contexts, or to a continuous context space, is immediate in principle, since Eq. (5) generalizes to a softmax over any number of sub-models, but the empirical question of how many contexts the brain entertains at once, and at what cost, is open. Second, the simulation of Section 4.2 specified the pragmatic and epistemic values as time courses. A full implementation would compute them from a generative model of the task in the induced reality, using the discrete-state formulation of active inference [12, 54, 151], and would let learning change the model over sessions. This is the natural next step, and it would allow quantitative predictions for the graded violation and epistemic value paradigms proposed above. Third, we have said little about the neuronal implementation. The hierarchical message passing of active inference has been mapped onto cortical microcircuits and onto the thalamo-cortical regulation of precision [56, 57, 65, 152], and the context inference of Eq. (5) presumably involves the same medial prefrontal and hippocampal circuits that support scene construction and reality monitoring [81, 153], but a specific proposal must await data from paradigms of the kind we have outlined. Fourth, the psychophysiological markers we have listed are correlates of precision and prediction error in general, not of the context inference in particular; their specificity must be established by co-registration with the events of both realities, as we argued in the two-competitor model. Fifth, individual differences, which we have located in the mantle [5], enter the account through the learned precisions and the prior preferences and are therefore expected to be large; the account predicts, for instance, that individuals with high interoceptive precision are less immersible by exteroceptive means and more immersible by embodiment, a prediction that can be tested. Finally, we have restricted ourselves to a single agent. Social presence, the sense of being with another person, which Abrash regards as the decisive application of immersive technology [9], requires a model in which the other agent’s states are inferred and in which the induced reality must support the sensorimotor contingencies of interaction, gaze, turn-taking, and touch; the framework extends to this case, but we have not developed it here.

## 7 Conclusions

We have proposed that immersion is inference. An induced reality is a candidate generative model that competes with the model of the actual reality for the explanation of a shared sensorium and for the control of action; the degree of immersion is the posterior probability that the induced reality is the current context; presence is the transparency of the winning model; and a break in immersion is a Bayesian switch driven by a high-precision prediction error that the induced model cannot absorb. The exogenous and endogenous components of the two-competitor model are two routes of precision allocation, one set by the evolutionarily entrenched core of the perceptual system and one set by the expected free energy of the agent’s policies, and the affective and motivational dynamics that the two-competitor model captured with a fatigue module are the pragmatic and epistemic value of remaining in the induced reality. The account recovers the quantities of the two-competitor model, reproduces its scenarios, and adds a perceptual variable, the context posterior, that the attentional focus vector drives but does not equal. This is the formal content of the hypothesis that attention is necessary but not sufficient for immersion: attention sets the evidence, and the expected free energy of the policies in the induced reality decides whether the evidence becomes belief. The account connects presence, embodiment, cybersickness, and narrative absorption to a single quantity; it specifies what the psychophysiological markers of immersion measure; and it yields a design hierarchy in which the core of perception is non-negotiable and the mantle can be trained. The engineering intuition that a virtual reality only has to satisfy the senses, not replicate the world, is thereby given its neuroscientific form: the senses are satisfied when the world they are offered is the one that best explains what they sense and best serves what the person is trying to do.

## Author contributions

LR: Methodology, Investigation, Writing (original draft), Writing (review and editing). KLG: Writing (review and editing). JV: Conceptualization, Investigation, Methodology, Writing (review and editing). ALF: Conceptualization, Formal analysis, Investigation, Methodology, Writing (review and editing). FIC-S: Conceptualization, Formal analysis, Investigation, Methodology, Validation, Visualization, Writing (review and editing). DJS: Conceptualization, Formal analysis, Investigation, Methodology, Software, Visualization, Writing (original draft), Writing (review and editing), Funding acquisition, Supervision.

## Data and code availability

No empirical data were used. The Python code that generates Figures 2 and 3 and the Induced Reality Simulator, a browser application in HTML5/JavaScript that runs without installation and in which all parameters and events of the model can be varied, can be downloaded from https://cloud.hiz-saarland.de/s/S79mStn5TagwpTC. The package also contains the settings of the two scenarios of Section 4.2. We will make the code publicly available on GitHub after acceptance.

## A Simulation details

The simulation of Section 4.2 uses a discrete time step of Δ*t* = 1 s over 20 min, that is, *T* = 1200 steps. For each reality *c* ∈ {ar, ir}, the pragmatic value *r*_*c*_(*t*) and the epistemic value *ε*_*c*_(*t*) of the policies acting in *c* are specified as time courses (Figure 2, first row), and the expected free energy of these policies is calculated as *G*_*c*_(*t*) = −((*r*_*c*_(*t*) + *ε*_*c*_(*t*)). The policy posterior is

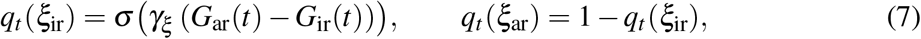

with *γ*_*ξ*_ = 4. Endogenous precisions follow Eq. (6) with modeling assumptions *ζ*_0_ = 0.35 and *ζ*_max_ = 1. Exogenous precisions 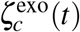 are oscillatory functions with a baseline of 0.5 (ir) and 0.45 (ar); the slamming door adds, from *t* = 10 min on, a causal alpha-function transient *xe*^1−*x*^ with *x* = (*t* −*t*_door_)*/*4 s (peak after 4 s) to 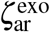 and an exponentially decaying epistemic value of amplitude 0.4 and time constant 35 s to *ε*_ar_; the brief transient and the slower decay of the epistemic value are what produce the fast break and the slower recovery in Figure 2, together with the precision-dependent update rate below. The precision-weighted sensory evidence of each reality is 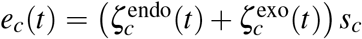, where *s*_*c*_ is the coherence of the reality’s model with its streams once attended (*s*_ir_ = 0.90, *s*_ar_ = 0.95). The context log-odds relax toward a target that combines the evidence differential with a policy-informed prior,

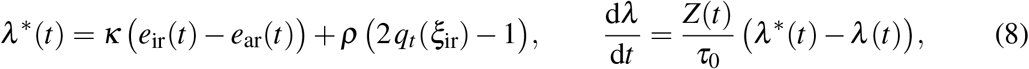

where *Z*(*t*) is the sum of all four precisions, so that the rate of belief updating scales with the total precision, and *κ* = 3, *ρ* = 1, *τ*_0_ = 15 s. The degree of immersion is *q*_*t*_(*c* = ir) = *σ* (*λ* (*t*)). The two-competitor projections are computed from the same precisions with the original transfer functions and parameters, 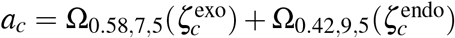, and *ϕ*_*a*_ = tan^−1^(*a*_ir_*/a*_ar_).

The free energy proxy is

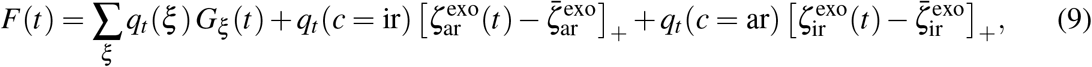

where [·]_+_ denotes rectification and 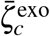 is the maximum of the expected oscillatory exogenous precision (0.53 for ar, 0.64 for ir), so that only unexpected exogenous events in the reality one does not currently believe in contribute unexplained prediction error. Valence is −d*F/*d*t* smoothed with a 15 s moving average. All quantities are dimensionless except time; the log-odds are initialized at *λ* (0) = 1.5, valence is expressed per minute, and the epistemic values have small floors (0.03 for ar, 0.02 for ir in scenario B). The pragmatic values *r*_*c*_ are affine proxies of the expected log-probability of preferred outcomes rather than log-probabilities themselves. The Python code is `simulate_immersion.py`, part of the code package given in the statement on data and code availability; the settings of scenarios A and B are included as `settings_preset_A.json` and `settings_preset_B.json` and can be loaded into the browser application.

